# Curating 16S rRNA databases enhances taxonomic accuracy and computational efficiency in microbial profiling

**DOI:** 10.1101/2025.11.04.686545

**Authors:** Mahdi Baghbanzadeh, Vedant Mahangade, Keith A. Crandall, Ali Rahnavard

**Affiliations:** Computational Biology Institute, Department of Biostatistics and Bioinformatics, Milken Institute School of Public Health, The George Washington University, Washington, DC 20052

**Keywords:** 16S rRNA gene sequencing, Reference databases, Database curation, Taxonomic classification, DADA2 pipeline, Amplicon sequence variants (ASVs)

## Abstract

The 16S rRNA gene serves as the gold standard molecular marker for microbial profiling, yet taxonomic assignment accuracy depends critically on reference database quality. Substantial heterogeneity exists among databases in sequence coverage, curation standards, and taxonomic nomenclature, leading to conflicting taxonomic assignments. Despite previous comparisons highlighting performance differences, the impact of database preprocessing, including sequence cleaning and redundancy removal, on taxonomic classification remains understudied. To improve database quality, we implemented novel cleaning approaches to remove nested sequences, duplicate sequences, and correct missing taxonomic nomenclature. We compared four major 16S databases (SILVA, Greengenes2, RefSeq, and MIMt) using 69 mock communities and the DADA2 analysis pipeline for microbial genus-level profiling. Database size was significantly reduced after cleaning: SILVA reduced from 452,055 to 291,733 sequences, Greengenes2 from 337,506 to 277,982 sequences, MIMt from 48,749 to 34,734 sequences, and RefSeq from 27,376 to 25,970 sequences. Greengenes2, MIMt, and RefSeq, which exhibited comparable performance, consistently outperformed SILVA in recall, precision, and abundance estimation accuracy across all sample types. Cleaning SILVA improved computational efficiency by up to 50% while maintaining classification performance. We provide a benchmarking framework with the cleaned databases as resources for accurate 16S rRNA analysis and profiling.

## Introduction

The 16S rRNA gene is widely used as a fundamental molecular marker in microbial ecology, evolutionary biology, and taxonomy due to its universal presence across bacteria and archaea and highly conserved regions interspersed with variable domains^1,2^. Using 16S rRNA as a marker gene enables researchers to identify and classify microorganisms within complex communities across environmental microbiology, clinical diagnostics, and microbiome research^3–5^. High-throughput sequencing technologies have enabled rapid, large-scale 16S analysis and standardization of sequencing workflows^5,6^. Despite limitations in species-level resolution and intragenomic heterogeneity, 16S rRNA gene sequencing remains the gold standard for microbial profiling^4,6,7^.

Amplicon sequence variants (ASVs) have emerged as a high-resolution alternative to operational taxonomic units (OTUs), enabling discrimination of sequences differing by a single nucleotide and improving reproducibility across studies^8,9^. However, ASVs can artificially split bacterial genomes due to intragenomic variation, requiring careful interpretation^10^. The DADA2 pipeline has become standard for robust error correction and taxonomic assignment^8^, comparing ASVs against curated reference databases like Greengenes2^11^, SILVA^12^, MIMt^13^, or RefSeq^14^ for taxonomic classification from phylum to species level^8,12–15^.

Taxonomic assignment accuracy is fundamentally dependent on reference database quality^16,17^. However, substantial heterogeneity exists among databases in sequence coverage, curation standards, and taxonomic nomenclature, with some being manually curated while others rely on automated processes^16,18,19^. These differences in annotation accuracy, redundant entries, and classification schemes result in conflicting assignments, making database choice critical for taxonomic classification performance^16,18–20^.

The current database landscape is shaped by distinct curation philosophies: Greengenes2 being driven by whole genome phylogeny^11^; SILVA offers community-curated comprehensive coverage with phylogenetic tree-guided manual curation^21^; RefSeq emphasizes high curation standards through NCBI’s Targeted Loci Project with rigorously validated sequences^14^; and MIMt^13^ adopts a hybrid approach integrating RefSeq and GenBank sequences with manual curation at all taxonomic levels, resulting in a compact yet highly accurate database excelling in species-level identification^13^. These databases represent the spectrum from broad community-driven curation to high-stringency institutional resources to innovative hybrid approaches to accumulate 16S data for computational analyses.

Previous database comparisons have highlighted substantial differences in coverage, resolution, and performance, with database choice significantly influencing microbial community analyses^17,20,22,23^. However, these studies lack investigation of database preprocessing effects, specifically cleaning sequence names and removing redundant sequences before comparison on mock communities.

In this study, we evaluated reference database choice impact on 16S rRNA profiling using 69 mock communities with known compositions and using four major databases: Greengenes2, SILVA, RefSeq, and MIMt. Each database was curated by updating taxonomic names, removing incomplete profiles, and eliminating duplicate and nested sequences, to ensure fair comparisons. Mock community reads were preprocessed using DADA2, with taxonomic profiling performed separately for each database while keeping all other variables constant. We assessed classification performance, computational time, and abundance estimation accuracy. Our results show Greengenes2, MIMt and RefSeq generally outperform SILVA, with SILVA exhibiting highest sequence redundancy and lowest coverage of distinct genera. Notably, after cleaning, SILVA achieved comparable performance in half the computational time. We also provide an automated pipeline for cleaning 16S databases, along with curated versions available at https://github.com/omicsEye/16S_DB.

## Methods

### Mock communities

We selected 69 mock communities previously established^24^ to evaluate database performance. These communities are categorized into four groups: “BEI balanced”, “BEI unbalanced”, “VAIO,” and “Simple”. The BEI samples contain reads from 17 distinct genera and families, while VAIO samples include reads from 13 distinct genera and families. The “Simple” group, a subset of VAIO samples, contains only one to five taxa. Detailed sample descriptions are available in ^24^.

### Sequence processing

We processed all raw FASTQ files using the DADA2 pipeline^9^ following standard quality filtering and denoising procedures. After preprocessing, we assigned taxonomy to the resulting amplicon sequence variants (ASVs) using the same DADA2 pipeline. All parameters remained constant throughout the analysis, with only the reference database varying among comparisons.

### Database selection and preparation

Four major 16S rRNA gene reference databases were evaluated in this study: SILVA 138.2 (preprocessed by DADA2 maintainers; available at https://zenodo.org/records/14169026), RefSeq 16S database of archaea and bacteria from the NCBI targeted loci project^14^, the MIMt 16S database^13^, which combines RefSeq targeted loci and GenBank sequences, and Greengenes2 (preprocessed by DADA2 maintainers; available at https://zenodo.org/records/14169078). To ensure compatibility with the DADA2 pipeline, we standardized the sequence headers in the RefSeq and MIMt databases using custom Python scripts and the seqkit command-line tool^25^.

A comprehensive database cleaning procedure was implemented to remove redundant and repeated sequences across all three databases. Duplicate sequences were identified and removed using the `seqkit rmdup` function, which detects exact matches between sequences while preserving only one representative sequence. During this process, detailed logs of the sequences and their associated identifiers and taxonomic names were maintained in separate files. Following duplicate removal, we identified nested sequences within each genus by first unifying taxonomic nomenclature and then extracting all sequences belonging to individual genera into separate files to minimize computational overhead. Within each genus-specific file, nested sequences were identified using the `seqkit locate` function. A nested sequence was defined as a sequence that was completely contained within another sequence (no mismatch or gap) from the same genus. Comprehensive logs of all identified nested sequences were maintained for reference. For all the databases, we only included the sequences that have genus-level information.

The curation processes resulted in four additional databases: GG2_c, MIMt_c, RefSeq_c, and SILVA_c, where the “_c” suffix denotes the curated version with duplicate and nested sequences removed. For SILVA_c, taxonomic profiles were also updated as described above.

### Performance evaluation

We assessed database performance using three standard metrics in taxonomic classification: recall, precision, and F1-score. Recall was calculated as the proportion of expected taxa that were successfully detected in the samples, representing the database’s sensitivity. Precision was determined as the proportion of detected taxa that were correctly identified, indicating the database’s specificity. The F1-score, calculated as the harmonic mean of precision and recall, provided a balanced measure of overall classification performance:

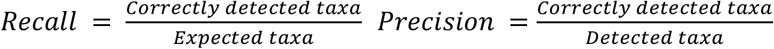

Additionally, normalized root mean square error (NRMSE) between the expected and observed relative abundances to quantify the accuracy of abundance estimations, previously published^20^:

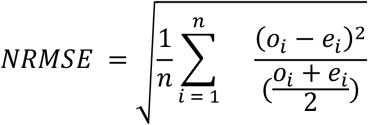

where for each sample with *n* taxa, *o*_*i*_ and *e*_*i*_ are the observed and relative abundance of taxon *i*.

## Results

### Database characteristics

The Greengenes2 database (GG2) initially contained 337,506 sequences. After removing 298 duplicated sequences and 12,261 nested sequences, the final Greengenes2 dataset comprised 277,982 sequences (GG2_c) covering 8108 genera and 2105 families (based on their specific nomenclature). Among these, there are 8,108 unique profiles up to genus level, including those with alphanumeric names such as “1-14-0-10-42-19”, “1XD8-76”. The MIMt database initially contained 48,749 sequences covering 4,951 genera and 897 families. After removing 13,623 duplicated sequences and 392 nested sequences, the final MIMt dataset comprised 34,734 sequences (MIMt_c). The SILVA database started with 452,055 sequences and 5,129 sequence IDs. Among these IDs, there were 4,063 unique complete taxonomic profiles up to the genus level, including those without an established name like “AB_125”, “LC_3”, and “*Candidatus Haloectosymbiotes*”, or cases like”*Escherichia-Shigella*”.

There were instances such as the whole ID for the sequence to be only as “*>Eukaryota*”, “*>Archaea*”, or “*>Bacteria*”. There were 40,658 duplicated sequences and 8,895 nested sequences. Our cleaning process resulted in a final SILVA dataset of 291,733 sequences (SILVA_c) covering 4056 genera and 650 families. The RefSeq database initially contained 27,376 sequences covering 4,255 genera and 785 families. After removing 384 duplicated sequences and 1,350 nested sequences, the final RefSeq dataset (RefSeq_c) contained 25,970 sequences.

### Taxonomic assignment performance

At the genus level, the MIMt database demonstrated superior performance compared to other reference databases examined in this study (**Figure 1a-e**). A comparison between the complete MIMt database and its curated version (MIMt_c) revealed comparable performance metrics in terms of recall and precision, with MIMt_c showing marginally higher averages (not statistically significant) in certain cases. Notably, since MIMt_c contains fewer sequences than the full MIMt database, it requires less computational time for taxonomic assignments (**Figure 1d**). Both MIMt and MIMt_c databases yielded the lowest root mean square error (RMSE) values when compared to the expected relative abundance profiles across all tested databases (**Figure 1c**), indicating their higher accuracy in estimating community composition.

**Figure 1:**
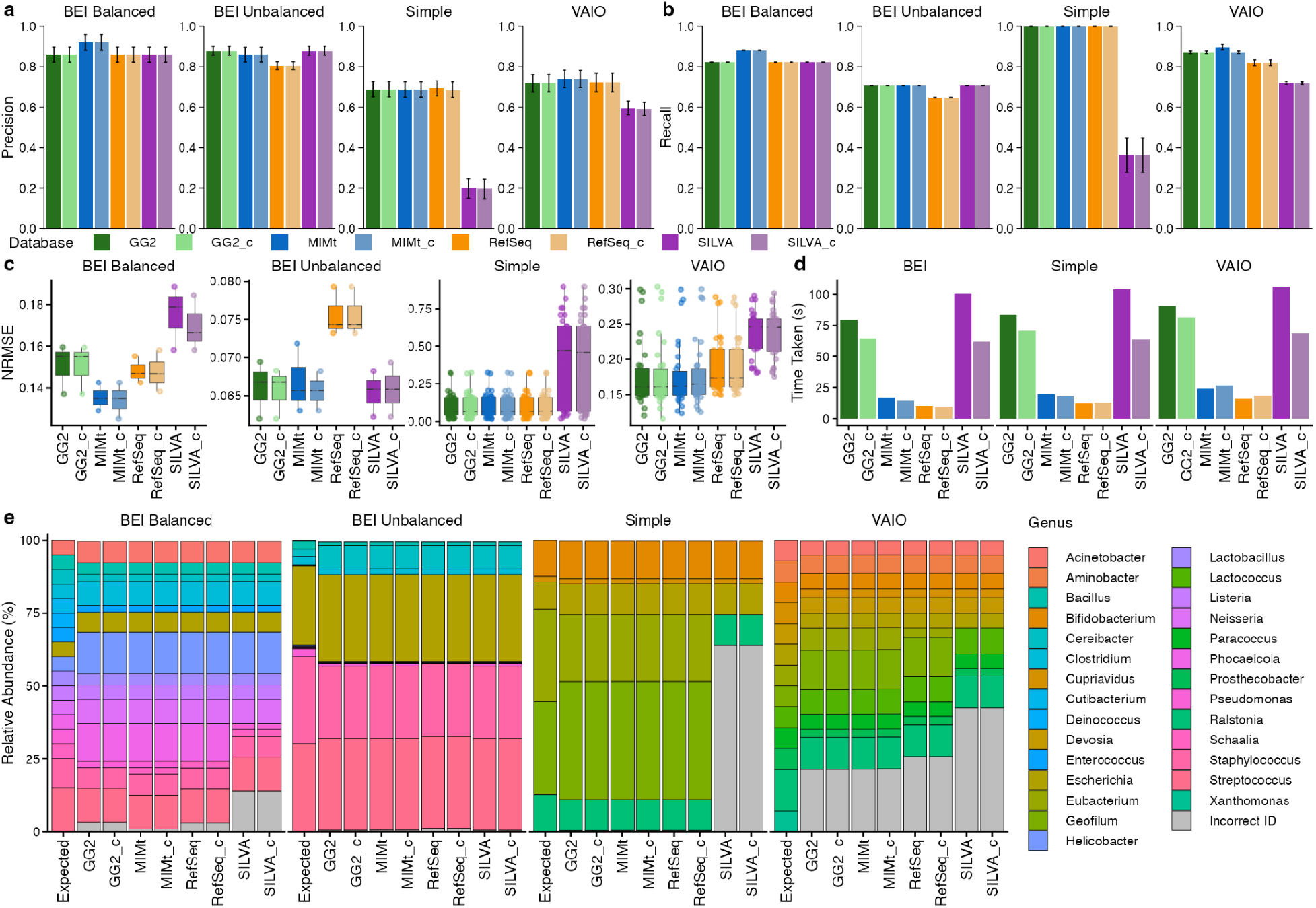
Comparison of DADA2 profiling performance using different 16S reference databases. **a**, precision of correctly detecting the genera in the samples. **b**, recall of correctly detecting the genera in the samples. **c**, NRMSE of relative abundances estimation by each database against the expected. **d**, computational time comparison. **e**, Relative abundances of genera within each type of sample.

The RefSeq and its curated counterpart (RefSeq_c) showed minimal differences in performance metrics. No substantial changes in recall and precision were observed between these two databases, except for the VAIO samples, where RefSeq_c demonstrated higher (not statistically significant) average recall scores (**Figure 1a-e**). The modest improvement in computational efficiency for RefSeq_c corresponds to the relatively small reduction in database size, with only approximately 5.1% of sequences removed during the curation process.

The curated SILVA database (SILVA_c) demonstrated a dramatic improvement in computational efficiency, requiring roughly 50% of the processing time compared to the complete SILVA database (**Figure 1d**). Despite this substantial reduction in database size and associated computational requirements, SILVA_c maintained equivalent performance levels to the complete SILVA database in terms of both recall and precision metrics (**Figure 1a-e**). Similarly, NRMSE values comparing taxonomic assignments to expected relative abundance profiles remained consistent between SILVA and SILVA_c (**Figure 1c**), suggesting that the curated database effectively preserved the taxonomic resolution capabilities of the full database while substantially reducing computational overhead. In the VAIO and Simple samples, SILVA and SILVA_c have a lower score of recall and precision from the results of other databases (**SUPPLEMENT 1**). Overall, the number of sequences in the database does not correlate with its performance.

Both SILVA databases exhibited notable taxonomic misclassification patterns that compromised their accuracy in microbial community profiling. Specifically, *Geofilum* remained undetected in both SILVA and SILVA_c analyses, while being successfully identified by all other databases tested, with SILVA incorrectly classifying these sequences as *Alkaliflexus* (**SUPPLEMENT 1**). This was also detected in another study^24^. Furthermore, *Escherichia* sequences are labeled as *Escherichia-Shigella* across all samples in both SILVA databases, maintaining identical relative abundances to the correctly identified *Escherichia* reported by other databases (**SUPPLEMENT 1**). Additionally, *Eubacterium* sequences were incorrectly classified as *Pseudoramibacter*, and *Phocaeicola* was reported as *Bacteroides* in SILVA-based analyses, representing a false identification that was absent from the original sample composition (**SUPPLEMENT 1**).

## Discussion

Our comprehensive evaluation of four major 16S rRNA gene reference databases reveals significant differences in taxonomic classification performance and computational efficiency. The SILVA database consistently performed worse than the others across all performance metrics due to misclassifying some of the samples, like *Geofilum* to *Alkaliflexus*. The MIMt database showed generally better performance than others, as it showed the lowest distance from the expected relative abundance, and this superior performance can be attributed to MIMt’s comprehensive sequence collection strategy, which combines RefSeq targeted loci with GenBank sequences, providing broader taxonomic coverage and more accurate genus-level classifications^13^. The database curation process proved highly effective in improving computational efficiency without compromising classification accuracy. SILVA_c achieved the most dramatic improvement, reducing processing time by over 50% while maintaining equivalent taxonomic resolution capabilities. The systematic misclassifications observed in SILVA databases highlight the effect of database selection and the critical limitations in current taxonomic nomenclature systems^16,26^. The misidentification of *Geofilum* as *Alkaliflexus*, confirmed by previous studies^24^, and the misclassification of *Phocaeicola* as *Bacteroides* and *Eubacterium* as *Pseudoramibacter* underscores the need for regular database updates to incorporate evolving bacterial taxonomy.

These findings have important implications for microbiome research accuracy and reproducibility. The choice of reference database can significantly impact community composition estimates and downstream analyses, particularly in studies focusing on specific bacterial genera^26,27^. Our automated and reproducible pipeline, along with its curated database, can be readily used and expanded as more data becomes available.

## Data and Code Availability

All data, codes, and processed databases are available at https://github.com/omicsEye/16S_DB.

## Acknowledgements

This work was supported by grants from the National Science Foundation (DEB-2109688; TI-2507498) and a Technology Maturation Award from George Washington University’s Technology Commercialization Office.

## Author contributions

M.B. developed the cleaning pipeline, performed benchmarking, and analyzed the data. V.M. assisted with testing, validation, and code refactoring. K.A.C. provided biological insights and contributed to the study design. A.R. conceived and supervised the project and guided the analyses. All authors contributed to writing and reviewing the manuscript.

## Competing interests

A.R. and K.A.C. are inventors of a pending patent related to the work presented and co-founders of seqSight, a company associated with the technology. The other authors declare no competing interests.

## Notes

https://github.com/omicsEye/16S_DB

